# The Structure of Oxalate Decarboxylase at its Active pH

**DOI:** 10.1101/426874

**Authors:** M. J. Burg, J. L. Goodsell, U. T. Twahir, S. D. Bruner, A. Angerhofer

## Abstract

Oxalate decarboxylase catalyzes the redox-neutral unimolecular disproportionation reaction of oxalic acid. The pH maximum for catalysis is ~4.0 and activity is negligible above pH7. Here we report on the first crystal structure of the enzyme in its active pH range at pH4.6, and at a resolution of 1.45 Å, the highest to date. The fundamental tertiary and quaternary structure of the enzyme does not change with pH. However, the low pH crystals are heterogeneous containing both a closed and open conformation of a flexible loop region which gates access to the N-terminal active site cavity. Residue E162 in the closed conformation points away from the active-site Mn ion owing to the coordination of a buffer molecule, acetate. Since the quaternary structure of the enzyme appears unaffected by pH many conclusions drawn from the structures taken at high pH remain valid. Density functional theory calculations of the possible binding modes of oxalate to the N-terminal Mn ion demonstrate that both mono- and bi-dentate coordination modes are possible in the closed conformation with an energetic preference for the bidentate binding mode. The simulations suggest that R92 plays an important role as a guide for positioning the substrate in its catalytically competent orientation. A strong hydrogen bond is seen between the bi-dentate bound substrate and E101, one of the coordinating ligands for the N-terminal Mn ion. This suggests a more direct role of E101 as a transient base during the first step of catalysis.

## 1. Introduction

Oxalate Decarboxylase (OxDC) is a Mn-dependent enzyme belonging to the cupin superfamily which is found in fungi and soil bacteria (1-5). The best characterized isozyme from *Bacillus subtilis* can be conveniently overexpressed in *Escherichia coli* (6). It catalyzes the heterolytic cleavage of the relatively inert carbon-carbon bond of the oxalate mono-anion, a unimolecular disproportionation reaction which is nominally redox-neutral. Yet, the presence of dioxygen is obligatory for catalysis and O_2_ is generally considered to act as a co-catalyst (6). Several crystal structures of the enzyme have appeared in the literature over the past 16 years all of which refer to enzyme poised at a pH of 7.5 or higher (7-14). Yet, OxDC is only active at low pH with peak activity at or below pH4 and negligible activity above pH7 (15). OxDC crystallizes as an aggregate of six monomers (see supporting information, fig. S1, for an illustration) (7). This homo-hexamer is likely also the active form of the enzyme in solution (6). The monomeric subunit weighs around 45kDa and contains two Mn ions labeled N- and C-terminal based on their proximity to the respective chain ends. Each of them is bound in two neighboring cupin domains by three histidine and one glutamate residues in approximate octahedral coordination leaving free valences for bound water and/or substrate. The function of the C-terminal Mn is unknown. However, removal of the C-terminal Mn or replacement with other metal ions leads to inactive enzyme (16).

The observation of two different conformations of a loop segment near the N-terminal cupin domain consisting of amino acids SENST161-165 has pointed to the N-terminal Mn binding pocket as the active site for OxDC (7-10). The open conformation features a solvent channel allowing substrate access to the Mn ion while the closed conformation can protect intermediates from leaking out, preventing interruption of the catalytic cycle. Based on kinetic isotope effect studies, a catalytic mechanism was proposed that involves radical intermediates with the surprising conjecture that dioxygen bound to the active-site Mn ion serves as a transient storage container for the electron removed from oxalate in an initial proton-coupled electron transfer (PCET) step (17). Evidence for the intermediate carbon dioxide radical anion, derived from oxalate was obtained by EPR spin trapping (10, 11, 15, 18). A superoxide radical adduct was also observed under turnover conditions (19). Interestingly, when one of the hinge residues of the flexible lid, T165, was replaced by valine the trapping yield for the carbon dioxide radical was increased by an order of magnitude while that for the superoxide radical decreased slightly (19). This observation may be taken as evidence that the two radicals emanate from different locations of the protein and that dioxygen might not actually bind at the active site. If dioxygen does not bind to the N-terminal Mn ion during catalysis, the substrate may have enough space to bind bi-dentate as one might expect oxalate to do.

Using a series of lid mutants, Burrell *et al*. concluded that E162 is essential for decarboxylase activity and suggested that it acts as the transient base that picks up the acidic proton from the oxalate monoanion during an initial PCET step (10). E162 turns inward toward the N-terminal Mn as seen in the closed-loop crystal structure by Just *et al*. (8) However, this conformation is unlikely to be the catalytically active one since it presents a steric clash with substrate bound to Mn (13). Saylor *et al*. reported on the structure of the T165V mutant enzyme which showed carbonate (or bicarbonate, a competitive inhibitor(20)) bound in the active site (11). As it turns out, either the T165V mutation or the inclusion of carbonate forces E162 into an orientation where its carboxylate oxygens point away from the active site. The fact that the T165V mutant exhibits a ten-fold lower decarboxylase activity compared to wild-type was interpreted as the result of this apparent ‘mis-folding’ of E162 into an unproductive conformation (11). More recently, Zhu *et al*. reported on the structure of a ΔE162 deletion mutant where Mn was substituted for Co (13). The deletion mutant contains oxalate in a mono-dentate end-on orientation which fits into the space vacated by the absence of E162. Based on modelling of the void space Zhu *et al*. rationalized that the original closed conformation of the SENST lid is only present in the absence of substrate or other small carboxylate ligands. They further argue that E162 may not function as a transient base in shuttling a proton back and forth but rather as a gating mechanism that locks the loop in the closed conformation during catalysis in order to prevent solvent access to the intermediate radicals (13).

In this contribution we report on the crystal structure of WT OxDC crystallized for the first time at a catalytically competent pH of 4.6 (PDB code 5VG3). It provides a clearer view of the catalytically competent structure of its active site and allows to validate conclusions drawn from earlier structural work. In the following, we will first focus on the asymmetric unit of the crystal and the quaternary structure of the enzyme. Critical connections linking the monomers in the hexameric cluster and explaining its stability are emphasized. We then turn our attention to the N-terminal Mn ion which shows a bound acetate molecule, and the flexible lid region that controls access to the active site. Specifically, we discuss the orientation of E162 in both the closed and open conformations at low pH. Furthermore, DFT (density functional theory) calculations are presented that probe the catalytically competent substrate binding mode demonstrating that both mono- and bi-dentate substrate coordination geometries are possible with the side-ways bi-dentate binding mode being energetically preferred. Finally, we take a look at the C-terminal site of the monomer where we observed acetate bound in a protein cavity with a narrow channel leading to the second Mn ion.

## 2. Materials and Methods

### 2.1 Expression and Purification of *Bacillus subtilis* OxDC

Expression and purification of recombinant His_6_-tagged WT OxDC were carried out following previously published procedures (8, 9, 11, 12). To remove dissolved metals from the preparation, Chelex 100 resin (Bio-Rad, Hercules CA) was added to the enzyme after the serial dialysis steps. The solution was shaken on ice for approximately one hour followed by removal of the resin. The protein solution was then concentrated using Amicon Centriprep YM-30 centrifugal filter units (EMD Millipore, Billerica, MA). Protein was further purified by FPLC using a HiTrap Q HP column followed by gel filtration on Superdex 200 10/300 GL column (GE Healthcare Life Sciences). OxDC was concentrated to 2 to 4 mg/mL for use in crystal screens. Protein concentration was measured utilizing a Bradford assay. ICP-MS determination of metal content was performed at the University of Georgia Center for Applied Isotope Studies Chemical Analysis Laboratory (Athens, GA). Typical batches were found to contain 1.6 Mn per monomer. Michaelis-Menten parameters of the decarboxylase activity of OxDC were determined by an end-point assay measuring the production of formate, as previously described (6, 8, 12). Typical values of *k*_cat_ ≈ 50 − 60 s^−1^ and *K*_M_ ≈ 1 − 4 mM are in line with literature data.

### 2.2 OxDC crystallization and structure solution

Initial crystallization conditions for native OxDC were tested using the sparse matrix screens: Crystal Screen HT (Hampton Research), using 2-4 mg/mL of purified protein. Optimized conditions were found to be 0.1 M sodium acetate trihydrate pH 4.6, 0.2 M sodium chloride, and 30% v/v 2-methyl-2,4-pentanediol. Diffraction quality crystals grew in 3 months using the hanging drop method at room temperature.

Initial data was collected on a Rigaku generator with R-AXIS VII++ detector and the data was processed using XDS and solved in PHENIX with PHASER (21, 22). The phasing model utilized was PDB ID 1UW8 (8). High resolution data was collected at Argonne National Laboratory on beam line 21-ID. Before data collection on the APS, crystals were soaked in mother liquor with 10 mM oxalate, however, the density maps did not reveal bound oxalate. Refinement was carried out with PHENIX-and iterative model building conducted in COOT (23, 24). Details of the data collection and refinement parameters are given in Table 1.

**Table 1:**
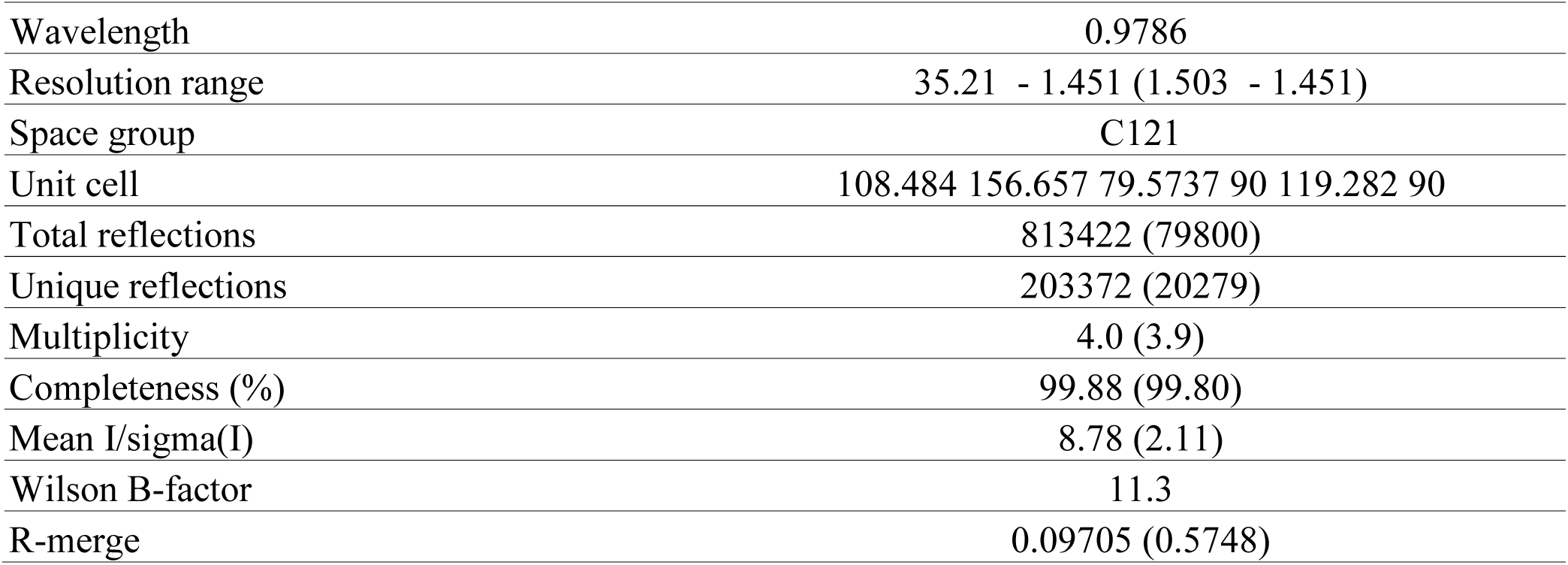
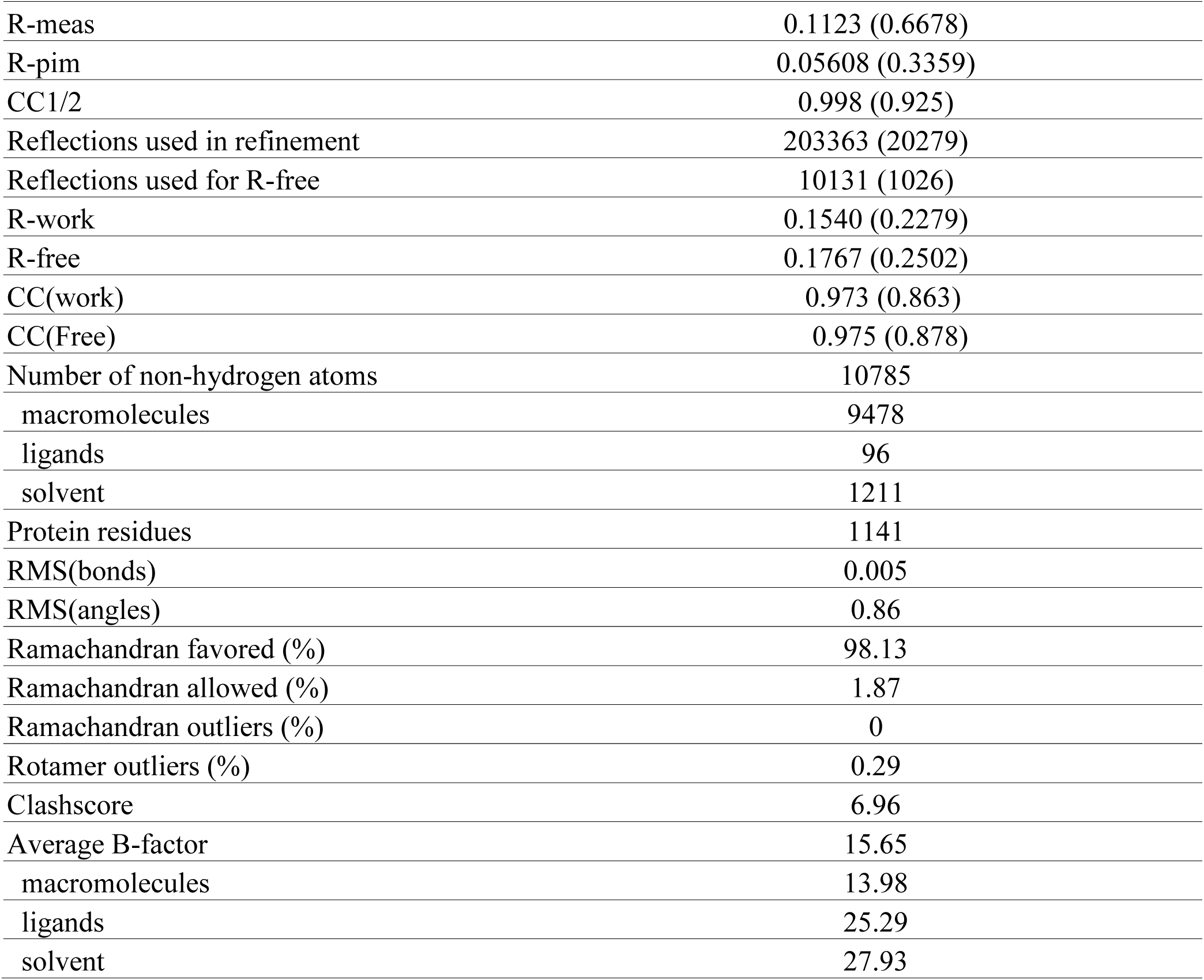
Data Collection and Refinement Statistics. Statistics for the highest resolution shell are shown in parentheses.

### 2.3 DFT Calculations

Oxalate was modeled in the active site using Gaussian09 (25). Active site residues were extracted and the backbone removed except for the α-carbons which were methyl-capped (26, 27). Both α- and β-carbons were frozen in place in order to simulate the protein environment. Residue R92 was included because of its experimentally observed impact on catalytic activity (9). Geometry optimizations were then performed using the CAM-B3LYP(28) functional and cc-pVDZ(29) basis set. Frequency calculations were performed at the cc-pVDZ level to ensure that a true minimum is reached. Following optimization, RMSD alignment was done with the crystal structure to orient the bound molecules using PyMOL (30, 31).

## 3. Results and Discussion

### 3.1 Quaternary Structure

The asymmetric unit consists of a trimer of OxDC forming an equilateral triangle linking two subunits at the vertices (supporting information fig. S1A). The biological unit of OxDC at low pH remains a homo-hexamer where two triangles stack face to face forming a dimer of trimers similar to all previously reported high pH structures (7-13). Several reports in the literature mention that the enzyme occurs in clusters of at least hexameric size in solution suggesting that this aggregation state may be relevant for catalysis (6, 32). The topology is identical to what was reported for the high-pH structures,(7) demonstrating that lowering the pH does not affect the quaternary structure. This is significant since it validates previous mechanistic proposals (8, 9). It also shows that the observed pH dependence of enzyme activity(4, 15) is not due to any pH-dependent structural changes of the protein but is a feature of the catalytic mechanism, most likely the higher availability of Mn(III) at low pH (33).

While some of the more obvious protein-protein interactions within the hexamer have been pointed out before,(7) a comprehensive analysis has not been reported. We have therefore employed *in-silico* alanine scanning using the DrugScore^PPI^ web interface at http://www.dsppi.de (34). It applies a knowledge-based scoring method that allows to estimate the change in Gibbs free energy for a protein composed of multiple chains when any given amino acid in a chain of interest is replaced with alanine, *i.e*., its side-chain reduced to a methyl group (35, 36). 179 amino acids were detected in the interface region around monomer A where all other monomers were considered as potential binding partners (numeric DrugScore results are tabulated in supporting information). Fig. 1A-C shows the residues with a ΔΔ*G* value of 1.5 kcal/mol or higher in the alanine scan while the scores for all 179 amino acids involved in the various interfaces are plotted in part D. A hot spot region involving four tryptophans was observed where two monomers (labeled A and B) within the same triangle link up. W96 (near the N-terminal Mn) and W274 (near the C-terminal Mn) form a π-stacked dimer with an average distance of 3.5 Å between their almost parallel molecular planes as has been pointed out before(8) (see supporting information, fig. S2). They are located at the base of two interlocking claws (compare with fig. S1C in supporting information) and are part of two β-strands that make up the N- and C-terminal cupin domains, respectively (see fig. 1C). On the other hand, W171 with a ΔΔ*G* = 3.11 kcal/mol is part of one of three short α-helices that make up the head of the claw of monomer B. A corresponding symmetry-related tryptophan is found for monomer A, W348, with only a slightly lower ΔΔ*G* of 2.12 kcal/mol. The high density of tryptophans coupled with good protection from solvent access makes this tetra-W motif an excellent example of a canonical hot spot (37).

Stabilization of the face-to-face association of the two triangles to form the hexamer is facilitated for each monomer pair in large part by the cross-over region of their N-terminal domains, I11–V36, which hug the C-terminal cupin of the juxtaposed monomer as seen in figures 1A and B, and S1B (7). There are two apparent hot spots near the tip of this flap, I21 and Y64. Neither of them is solvent-exposed.

It is well known that ion pairs in proteins preferentially locate on the surface and if they are buried they require a polar environment or need to be clustered in order to be stabilizing (38). On the other hand, complex salt bridges with more than two acid/base partners help proteins assume their tertiary and quaternary structural organization necessary for proper folding and function (39, 40). Two network salt bridges contribute to the stabilization of the hexamer (see supporting information fig. S3 for illustration). They belong to a large class of so-called charge clustered salt bridges (41). The first one involves residues R290, E322, and K381 from monomer A as well as E178 from monomer B (see fig. S3A) and are found near the vertices of the triangles. The salt bridge is partially solvent exposed and shows several close-range distances that may contain hydrogen bonds in a network of four residues. The high-pH structures, 1J58 and 1UW8, show E178 swiveled about its C_α_−C_β_ and C_β_−C_γ_ bonds leaving only one potential hydrogen bond to R290 in place in an arc-shaped network (see fig. S3B). It appears to be the only protein-protein interaction motif in OxDC affected by the pH change. It tightens up into a more closely clustered topology at low pH perhaps to accommodate more protons with a larger number of polar connections.

**Figure 1:**
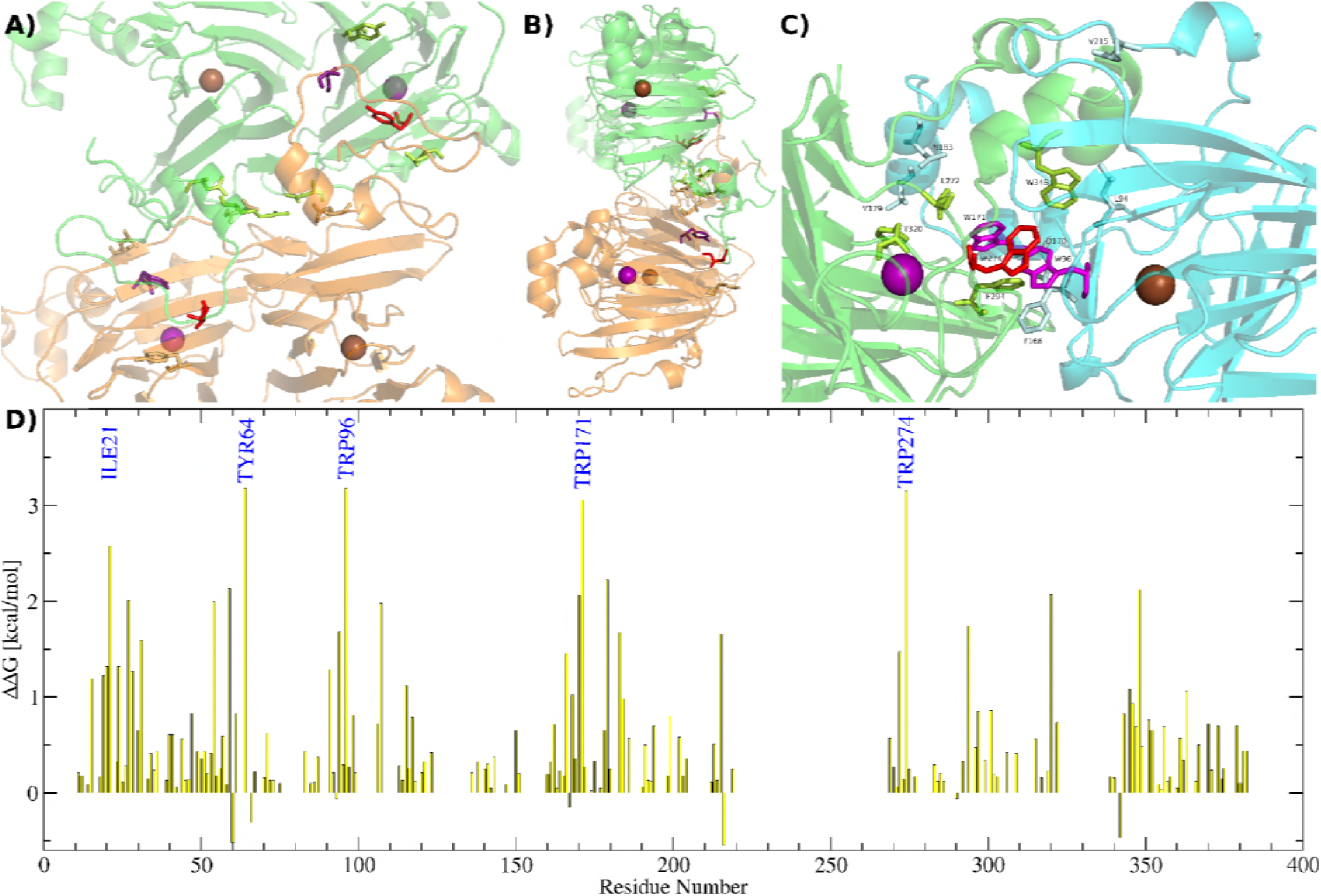
Results of in-silico alanine scanning of monomer A against all other monomers in the hexamer. Amino acids were classified as hot spots if their replacement by alanine resulted in a Gibbs free energy change of 2.5 kcal/mol or above (listed in part D). These residues are shown in panels A – C with red color if they belong to monomer A and in magenta if they belong to monomers B or D. Amino acids whose replacement with alanine results in a change between 1.5 and 2.5 kcal/mol are also shown. Those belonging to monomer A are shown in lime green while those belonging to monomers B and D are visualized in light cyan or orange, respectively. As a reference the N-terminal Mn ions are shown in purple, the C-terminal ones in brown. A) Visualization of the interface region between monomers A (green) and D (orange). Same view point as in figure 1B. B) Same as A) but viewed along the side of a triangle. C) Interface region between monomers A and B at the corner of a triangle. D) Results from the DrugScore^PPI^ calculation for 179 amino acids in the interface region of monomer A. Five amino acids were identified as being part of potential hot spots: I21, Y64, W96, W171, and W274.

The second networked salt bridge is found in the center between the short α-helical domains of the two flaps (center of fig. 1A). It forms between residues R27 and D54 from two adjacent monomers. These residues show large degrees of buriedness in the alanine scan of 9.13 and 11.35, respectively, and tie both monomers together at two different points on each in what looks like a cross stitch pattern (see fig. S3C and S3D). This motif is conserved in all published OxDC structures. Salt bridges with similar ‘quadrangle’ topology have been detected in a variety of different proteins where they participate in stabilizing the tertiary structure or in the formation of quaternary assemblies (42).

These protein-protein interactions stabilize the hexamer in solution and hold the key to understanding the organization of active protein. Disruption of these hotspots and salt bridges will be necessary in order to test for catalytic competency of smaller building blocks but is beyond the scope of this report.

### 3.2 Flexible Loop Region

During the refinement process we discovered that the crystals contained two alternate conformations of the enzyme that differed mainly in the position of the flexible SENST161-165 loop. We find an open conformation with an occupancy of between 40% and 50% which corresponds very well to the one reported by Anand *et al*. (7). A closed conformation with a weight of 50 to 60% is also found but differs from the one reported by Just *et al*. (8). The electron density map of this region of the protein from the X-ray structure is shown together with the best-fit models of both open and closed conformation in the supporting information, figure S4. Both conformations are compared with their high-pH counterparts in figures 2A and 2B. Interestingly, we find that acetate, a buffer molecule, is coordinated to the N-terminal Mn ion at low pH. The closed conformation observed by Just *et al*. (8) at high pH shows a similar backbone structure but differs significantly in the placement of the E162 residue. The dihedral angle along the E162 side chain (starting with the α-carbon) changes from 70.6° to −174.9° at low pH to avoid a steric clash with the acetate ligand (fig. 2B). In fact, the tucked-in conformation of E162 is only observed in OxDC structures that lack ligands bigger than water at the two free valences of the N-terminal Mn. Zhu *et al*. have simulated the void space in the high-pH closed structure, 1UW8, and noted that it would present a steric clash with the substrate even if oxalate was pushed further into the active side and coordinated to the Mn in a side-ways bidentate fashion (13). It is therefore safe to conclude that the high-pH closed conformation seen in 1UW8 is not a reasonable model of the active site under turnover conditions. A similar conformation to ours with E162 pointing away from the N-terminal Mn has been observed by Saylor *et al*. for the T165V mutant of OxDC (see comparison with T165V in supporting information, fig. S5) (11). It contains a bound carbonate or bicarbonate molecule which prevents E162 from pointing toward the Mn ion. Its conformation is therefore not a ‘mis-folded’ unproductive conformation as suggested before(11) but a necessity to provide enough space for the substrate and its reaction products. Acetate is a buffer molecule and is known as a weak competitive inhibitor of OxDC with an estimated *K_i_* of >100mM (20). Acetate binding has previously been observed at the N-terminal Mn(III) ion by EPR spectroscopy where it affects the magnetic fine structure parameters just like other small carboxylates, *i.e*., formate or succinate (33). Moreover, it has been suggested to be responsible for a decrease in the magnetic fine structure parameter of the N-terminal Mn(II) (43). It is therefore gratifying to finally see a crystal structure of OxDC with acetate bound.

Zhu *et al*. recently reported on the crystal structure of an E162 deletion mutant protein in which the Mn ions were substituted for Co, PDB code 5HI0 (13). The mutant does not turn over oxalate when Co replaces Mn. In fact, a molecule of oxalate was found coordinated end-on mono-dentate to the Co ion in the N-terminal site. Based on comparison with earlier structures the authors suggested that rather than functioning as a proton acceptor E162 acts as a gate for catalysis and may be necessary to lock the flexible loop into a catalytically competent conformation in the presence of substrate. It is therefore instructive to compare our low-pH closed structure with the deletion mutant OxDC-ΔE162. Figure 2C shows these two structures aligned with each other. The main difference is seen for residue N163 (N162 in the deletion mutant). It is pulled significantly toward the active site in ΔE162 because the backbone of the flexible SENST loop is stretched and loses its partial 3-10 helix motif as observed by DSSP (Define Secondary Structure of Proteins) analysis(44) (see fig. 2D). This leads to some instability resulting in much higher observed B factors of on average 61 Å^2^ in the ΔE162 OxDC variant compared to our structure (~11 Å^2^).

**Figure 2:**
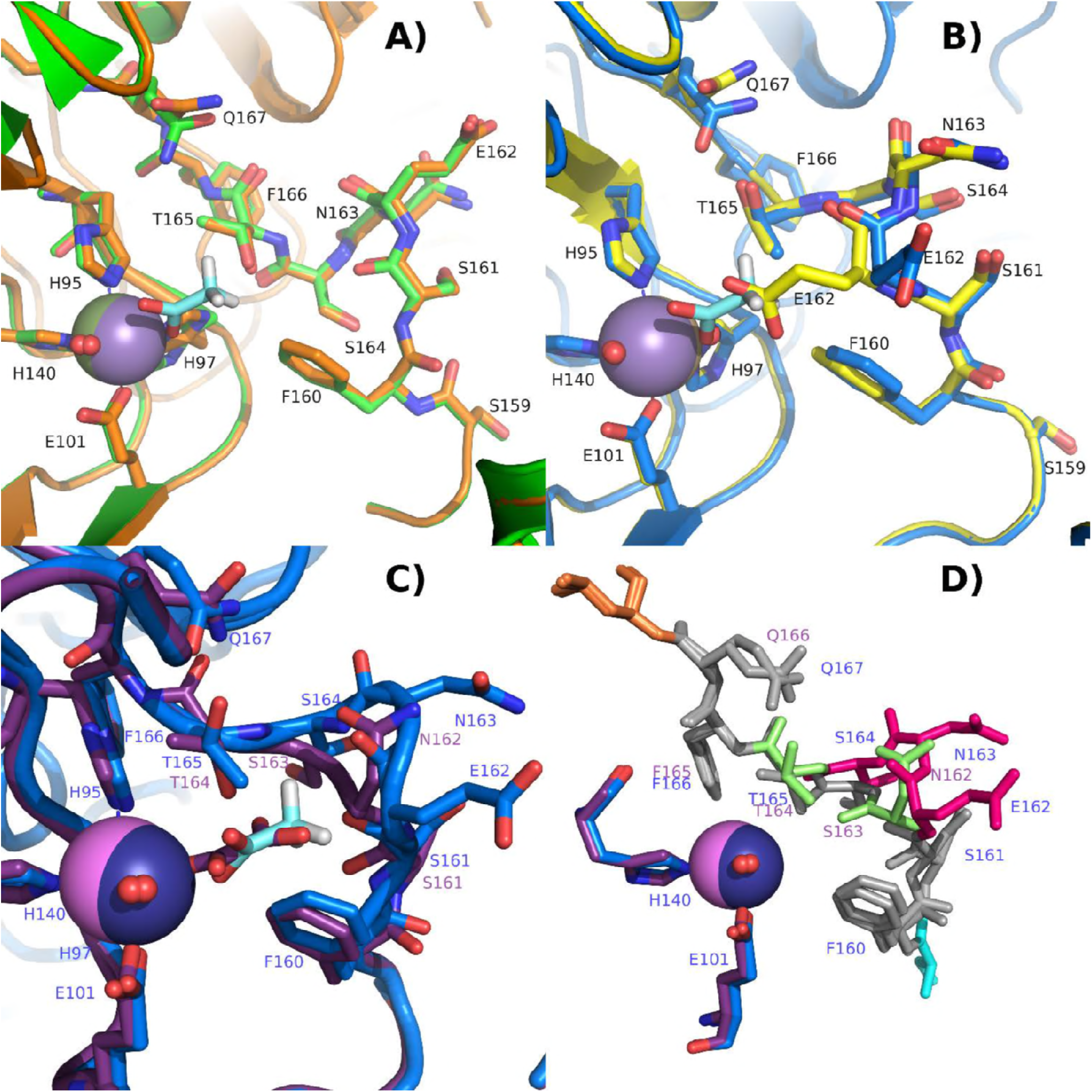
*A) Comparison of the open structure at high pH*(7) *(green) with the low pH open conformation (orange). B) Comparison of the closed structure at high pH*(8) *(yellow) with the low pH closed conformation (blue). Acetate in the low pH structure is shown with carbons in cyan. Its location would bring a steric clash with E162 in the high pH closed lid conformation. C) Overlay of the low pH closed conformation (blue) with the structure of a Co-substituted ΔE162 deletion mutant (purple). D) DSSP analysis of the same structures as in C). The colors reflect the color coding of the DSSP analysis. Red indicates a 3-10 helix, orange an α-helix, green a bend, cyan a turn, and grey no secondary structure assignment*.

The B factors of the two loop conformations at low pH show values comparable to the rest of the protein (see supporting information, fig. S6). Elevated B-factors in the coordinates of the flexible loop region have been noted before in the high-pH structures and were interpreted as indicative of the conformational flexibility of the lid (8). The fact that both conformations show significantly lower B-factors in our structure means that at low pH in the presence of a ligand the loop settles down in these two approximately equally stable and well defined conformations which exist in an equilibrium with approximately the same thermodynamic stability. This fact should in part be reflected in their respective H-bonding networks which prompted us to survey the polar bonding environment of the SENST loop and compare it with earlier analyses (8, 9, 11).

We focused on polar distances between 2.3 and 3.5 Å which are typical for intermediate H-bond lengths (45, 46). Figure S7 in supporting information indicates potential H-bonds between the polar parts of the SENST loop in the closed and open conformations. The main difference compared to the high-pH structures is again seen in the closed conformation since 1UW8 shows E162 tucked inside toward the Mn ion. This position of E162 is stabilized by H-bonds with R92 as well as two water molecules one of which is itself coordinated to the Mn ion. On the other hand, our low-pH structure shows no potential H-bond between E162 and surrounding residues which speaks against its role in stabilizing the closed conformation as Zhu *et al*. have suggested (13). At low pH the closed conformation is mainly stabilized by a rotation of the T165 side group away from its open-state position which allows it to strongly interact with Q167 within the same chain. It should be noted that the low pH structures show Q167 swiveled around its C_βγ_ axis with a change of dihedral angle from 166° to 71° which brings the amide group closer to T165 allowing for additional H-bond stabilization which appears to be absent at high pH (see fig. 2A and 2B). The open loop conformation shows fewer H-bonding interactions than the closed one. Yet, a close polar contact exists between E162 and H299 of the adjacent subunit along the hexamer axis with a heteroatom distance of only 2.6 Å, consistent with a well-established strong H-bond between these two residues as had been pointed out before (7-9). One additional interaction between S164 and the intra-chain E99 is noteworthy at the other end of the loop (9). The fact that residues from neighboring subunits participate significantly in the hydrogen bonding networks, particularly in the open conformation, suggests that the quaternary organization of the enzyme plays a role in catalysis. It is tempting to assign H299 the function of a ‘strike’ that holds on to the ‘latch bolt’ (E162) to keep the ‘door open’ to allow substrate to enter the channel. Indeed kinetic analysis of the mutant protein H299A revealed a ~50% loss of activity (9).

### 3.3 Substrate Binding

The closed conformation of 5VG3 so far provides the best structural model for the catalytically competent active site because of the pH at which the protein was crystallized and the presence of a carboxylate ligand where substrate is expected to bind. We were therefore interested in modeling the cavity by replacing acetate with oxalate in hopes that this might give clues about the binding mode of oxalate, *i.e*., mono-versus bi-dentate. Precedence for both binding modes exist. The Co-substituted deletion mutant ΔE162 described by Zhu *et al*. showed mono-dentate bound oxalate (13). On the other hand, a predicted oxalate decarboxylase (tm1287) from *Thermotoga maritima*, PDB #1OT4, shows a bi-dentate bound oxalate in a comparable sized binding pocket (14). A PDB data base search revealed that out of 49 metal-oxalate structures in proteins deposited in the data bank (oxalate is mostly coordinated to Mg^2+^ and Mn^2+^) 47 show bi-dentate binding geometry which is not surprising since oxalate will naturally bind side-ways bi-dentate if there is enough space available. This appears to be the case in OxDC and it is therefore plausible that once oxalate binds it rearranges to a bi-dentate coordination. We employed DFT calculations to further explore this possibility. Results of calculations for the mono- and bi-dentate binding mode of oxalate are shown in figures 3A and 3B, respectively.

As it turns out, both binding modes were well simulated and represent true energy minima. Both compare well with established crystal structures, the deletion mutant ΔE162 OxDC for the mono-dentate case, and the oxalate binding site of tm1287 from *Thermotoga maritima* for the bi-dentate case (see fig. S8 in supporting information). It should be noted that tm1287 simply serves as an example of a mono-nuclear Mn-containing protein with oxalate bound in a side-ways bidentate fashion. We do not wish to imply that oxalate is a substrate of this protein. Since the number of atoms and electrons was conserved in both models it was possible to compare total energies. As expected, the bi-dentate coordination of oxalate is lower in energy by about 28.5 kcal/mol. Of course this value does not take into account most of the second coordination shell and possible additional structural changes further away from the active site and is therefore only a qualitative indication of the well-known fact that oxalate prefers side-ways bi-dentate coordination to Mn(II).

**Figure 3:**
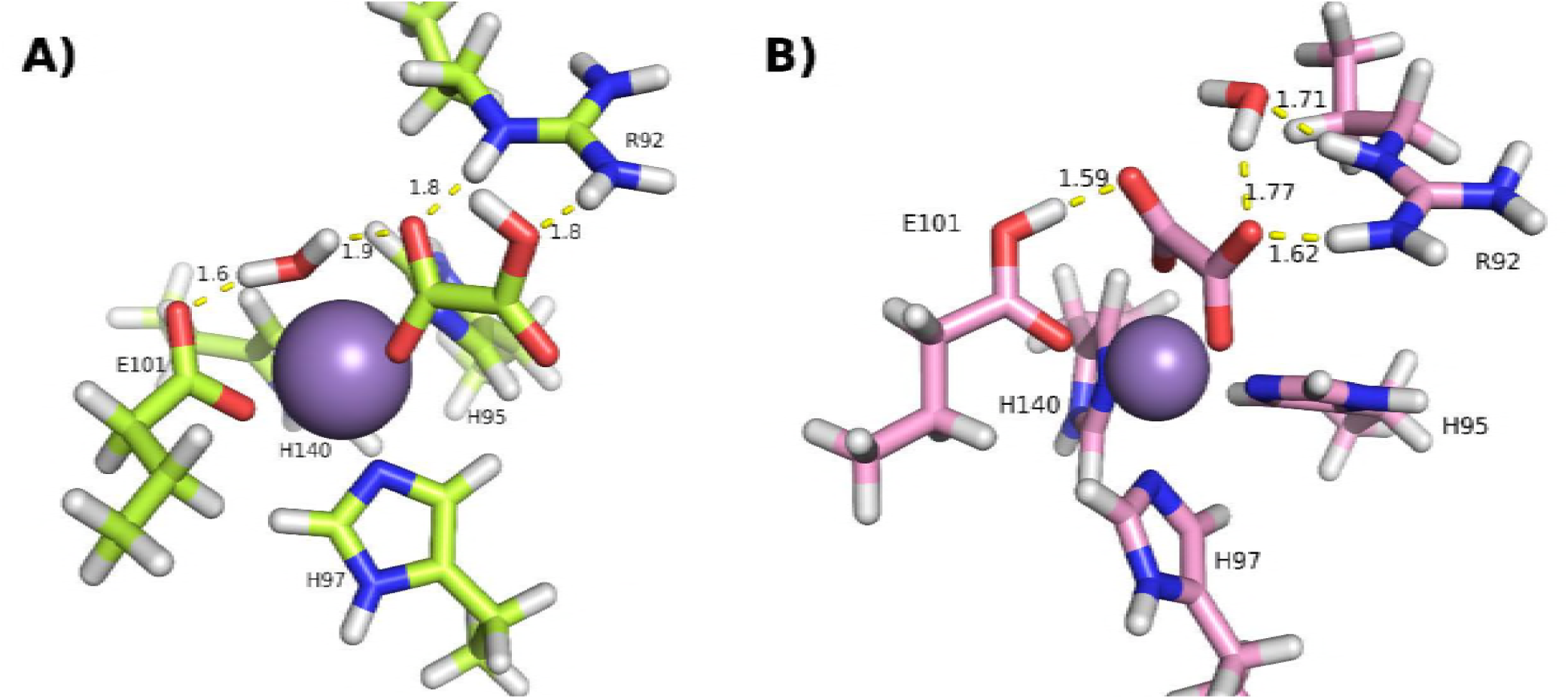
Results of DFT simulations of oxalate binding to the N-terminal Mn(II) ion. A) mono-dentate; B) bi-dentate. H-bonding distances between oxalate and its neighbors are shown with yellow dashes. The quoted numbers are in Å.

In the case of the mono-dentate model, oxalate binds in a similar orientation as acetate does in our low pH structure. It should be pointed out that R92 sits in close proximity to the oxalate ion with two of its amine nitrogens within hydrogen bonding distance to two oxygen atoms on oxalate. Water which makes up the sixth metal ligand is also part of this hydrogen bonding network as well as the free oxygen on E101. In the bidentate model oxalate is further twisted and establishes a potential H-bond with only one of its oxygen atoms to a primary amine of R92 since it is now shifted further into the active site pocket. The water molecule that was coordinated to Mn in the mono-dentate binding model now provides an additional hydrogen bond bridge between oxalate and the secondary amine group of R92. Interestingly, the hydrogen atom on the oxalate mono-anion has shifted to the other end of the molecule and in our simulation has transferred to the free oxygen on E101 leaving oxalate as a di-anion. This suggests that in addition to coordinating the N-terminal Mn ion E101 takes on the role of a transient base although this has yet to be experimentally confirmed. While our simulations do not yet allow to decide on the catalytically competent binding mode of the substrate they demonstrate that there is clearly enough space for either possibility (see also fig. S9 in supporting information). Earlier molecular simulations of a bi-dentate oxalate coordination by Just *et al*. also concluded that there was enough space available but raised doubts about this possibility because of the perceived lack of stabilizing hydrogen bonds and an unfavorable interaction between oxalate and I114, a residue in the second coordination shell of the N-terminal Mn ion (8). This earlier work used a molecular dynamics model rather than DFT and replaced the Mn with Fe since potentials for Mn were not readily available at the time. It is of course true that as oxalate inserts itself deeper into the N-terminal binding pocket it will interact with hydrophobic residues, particularly L153 and I142. Since we did not include these in our model small deviations in both energy minima and optimized geometries of substrate and surrounding residues are to be expected and still need to be clarified in more resource intensive future calculations that need to include the full second coordination shell. However, our model demonstrates several stabilizing hydrogen bonds to be present in the side-ways bi-dentate binding model. The proximal non-coordinating oxalate oxygen and E101 undergo a strong hydrogen bond while the distal non-coordinating oxalate oxygen is in H-bonding distance with one of the primary amines in R92 which moves toward the oxalate from its unperturbed position in the crystal (see figs. S8C and S8D). Moreover, our model shows a free water molecule nearby which had been replaced from coordinating the Mn ion by oxalate. It undergoes hydrogen bonding with both the oxalate and R92.

An interesting aspect of our simulations is the involvement of R92 in guiding oxalate into the active site and perhaps positioning it for catalysis. Site-directed mutagenesis of R92 has been performed before pointing to an essential role for this residue. R92L suppresses all activity and R92K severely diminishes it (8). Initially, we performed the geometry optimization of oxalate in the active site without including R92. However, the resulting position of oxalate in the mono-dentate coordination did not line up well with its structure in the deletion mutant. When R92 is included in the simulation it readily undergoes hydrogen-bonding interactions with the substrate. The dihedral angle of oxalate is somewhat larger in the bi-dentate conformation with 36.3 □ as compared to 21.0□ in the mono-dentate model. This can be explained by hydrogen bonding of both free oxygen atoms in opposite directions since the oxalate anion has a low rotational barrier and is well known to adjust its dihedral O-C-C-O angle to hydrogen bonding forces (47, 48). The close association between R92 and the substrate in our simulations is in agreement with the earlier mutagenesis results(8) and suggests a critical role for R92 to guide oxalate into place for catalysis. Hydrogen bonding between R92 and the mono-dentate bound oxalate in the structure of the ΔE162 deletion mutant was recognized before, yet the heteroatom distance was larger with 3.3 Å compared to the 2.8 Å observed in our model (13). It is well known that flexible loops are often indicative of induced-fit mechanisms of enzyme catalysis (49, 50). It would therefore not be surprising to find such a mechanism to be active in OxDC. It requires a trigger, *i.e*., a conformational change upon initial ligand binding, such as the motion of the R92 side-chain toward the substrate induced by the formation of strong hydrogen bonds. As a result of such a conformational change a destabilization of the open loop conformation may be expected resulting in the closing of the lid with subsequent repositioning of the substrate. It should be noted that electrostatic interactions between the positively charged R92 and the negatively charged substrate may also be crucial in bringing them into close contact during catalysis.

### 3.4 C-Terminal Site

We now turn our attention to the C-terminal Mn binding site. While it has been established that there needs to be Mn bound for activity,(16) its specific function is still unclear (51, 52). Based on EPR spin trapping experiments on the T165V mutant enzyme it was speculated that it could be the location where dioxygen binds and where superoxide is generated (19). It is therefore interesting to compare the conformation of the C-terminal Mn binding site of the low pH structure with the older structures at high pH. Figure 4A clearly shows that the amino acid coordination of the Mn ion is unchanged compared to both the open and closed structures at high pH (7, 8). Note that the low pH structure does not show any heterogeneity in the C-terminal site. It should be remembered that structure 1UW8 (closed N-terminal loop conformation) only shows one coordinating water ligand for the C-terminal Mn (8).

We noticed an acetate molecule bound in a cavity near the C-terminal site (see fig. 4B and 4C). It does not have access to the Mn-ion because residues R270 and E333 are part of the second coordination shell shielding the Mn against solvent access. However, the carboxylate oxygens of the acetate participate in an extended hydrogen bonding network involving both of these residues together with one of the two coordinating water molecules. The high pH crystal structures also show this cavity, but without bound acetate. There, the cavity appears to be somewhat smaller leading to a steric clash with acetate as seen when the structures are aligned (see supporting information fig. S10).

**Figure 4:**
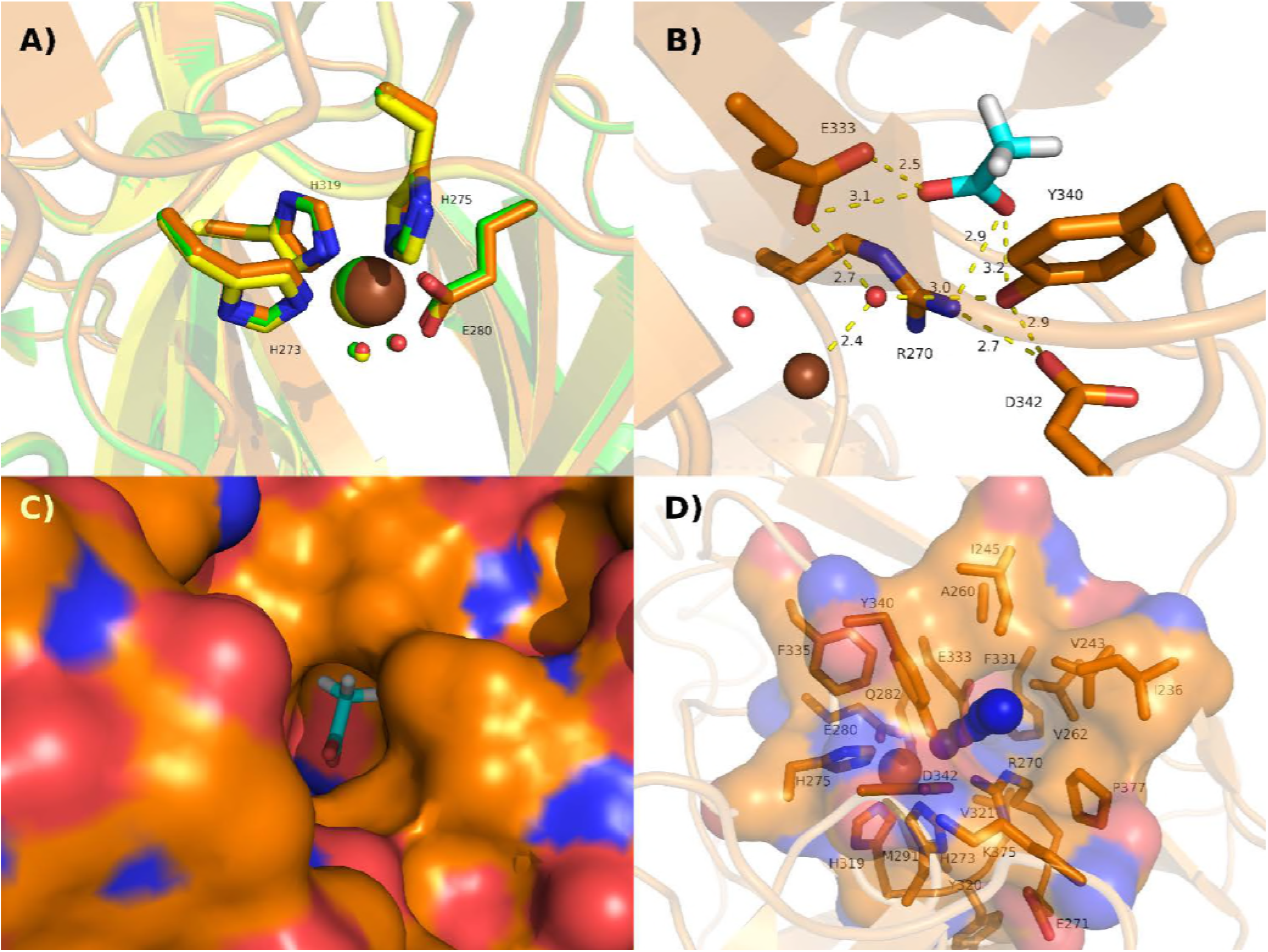
C-terminal Mn binding site. A) comparison of the Mn-coordination environment of the low pH structure) in orange with the original two high pH structures, 1J58(7) (green) and 1UW8(8) (yellow). B) Hydrogen bonding network between the acetate molecule bound in a cavity near the C-terminal Mn site and four nearby residues. C) Surface visualization of the protein with the cavity holding bound acetate. D) Similar view as in C) but acetate removed. A narrow channel is depicted with blue spheres that leads from the bottom of the cavity to the C-terminal Mn ion. The amino acids lining the channel are shown with orange sticks and are labeled. Their surface is indicated by the shaded area.

Finally, figure 4D shows a narrow channel that leads from the C-terminal Mn ion through the acetate binding site into the solution. This channel was simulated using the Caver 3.0.1 software package (53, 54). It has a static diameter of only 0.7 Å and is therefore too narrow for substrate. However, it could well be used by small molecules such as dioxygen or the superoxide anion. When acetate binds as seen in the low pH structure this channel is occluded.

## 4. Conclusions

The low pH structure is remarkably similar to the ones with enzyme at high pH. We conclude therefore that the pH dependence of catalysis is not based on structural changes of the protein. Moreover, the different position of E162 in the closed conformation is very likely not due to the change in pH but the presence of a small molecule ligand coordinated to the N-terminal Mn ion, acetate in our case. Since the carboxy group of E162 points away from the active site it is unlikely to serve as the transient base necessary for the rate-determining first step of catalysis, a PCET reaction.

Several key interactions between the OxDC monomers within the hexameric quaternary structure are critical for its stability in solution. A hydrophobic ‘hot spot’ containing four tryptophans and a networked salt bridge link the tips of two monomers together at the vertices of the triangles. Another, networked salt bridges involving residues R27 and D54 of two neighboring subunits are crucial for keeping the triangles together as a hexamer.

The low pH crystal contains both conformations of the SENST161-165 flexible loop in almost equal proportions suggesting that the closed and open loop have similar thermodynamic stability and are in equilibrium with each other, perhaps more so than at high pH.

DFT calculations allowed us to model substrate into the active site with two different coordination geometries. The side-ways bi-dentate conformation is spatially allowed and energetically preferred. R92 is seen to remain in hydrogen-bonding distance with the substrate in both cases and may function as a ‘guide’ to bring the substrate into its catalytically competent position. E101, one of the inner-sphere coordinating ligands, establishes a strong hydrogen bond with bidentate bound substrate and may take the role of a transient base in initial PCET reaction if this is the catalytically competent substrate binding geometry.

## Supporting Materials

Supporting materials containing ten figures and one data table are available.

## Author Contributions

MJB and UTT carried out the X-ray crystallography work under the guidance of SDB. JLG carried out the DFT calculations under the supervision of AA. AA designed the research and wrote the manuscript.

## Acknowledgments

This research was supported in part by the National Science Foundation, awards CHE-1213440 to AA and CHE-1411991 to SDB. JLG thanks the University of Florida for support through a Graduate School Fellowship.

---

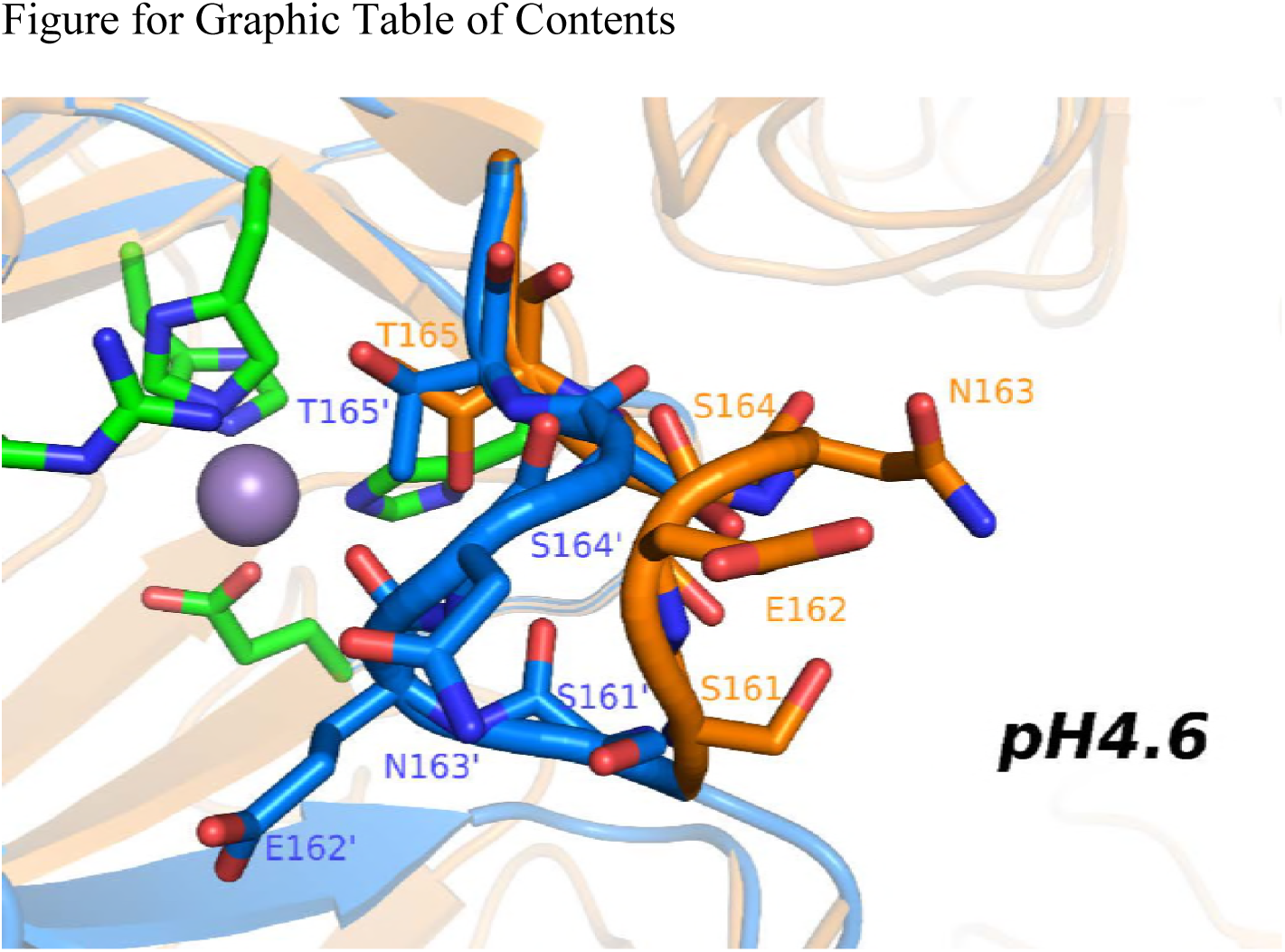

## The Structure of Oxalate Decarboxylase at its Active pH

Matthew J. Burg, Justin L. Goodsell, Umar T. Twahir, Steven D. Bruner, and Alexander Angerhofer, Department of Chemistry, The University of Florida, Box 117200, Gainesville, FL 32611, USA.

## Supporting Information

1. Quaternary structure of OxDC.

**Figure S 1:**
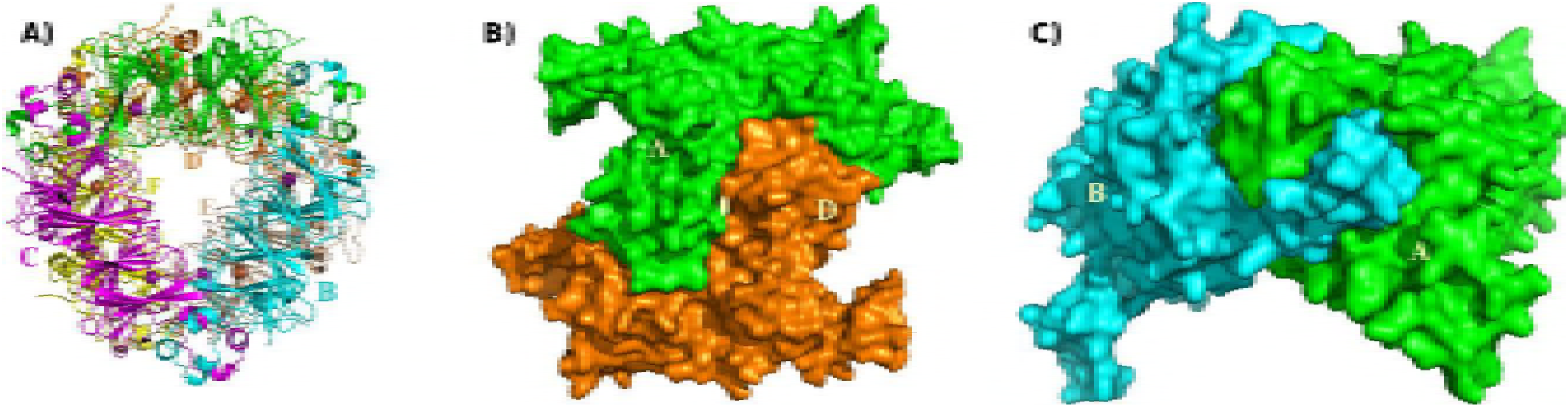
Quaternary structure of OxDC. (A) Top down view of the hexamer composed of a dimer of trimers. In clockwise orientation starting at the top with the triangle in the front we label the subunits A (green), B (cyan), C (magenta), D (orange), E (gray), and F (yellow). Mn ions are shown as purple (N-terminal) and brown (C-terminal) spheres. (B) Surface map side view of the hexameric structure seen from the outside of the hexamer and showing interlocking monomers A and D. (C) Corner of a trimer showing how the C-terminus of monomer A interlocks with the N-terminus of monomer B.

2. Hydrophobic hot spot near the vertices of the triangles.

**Figure S 2:**
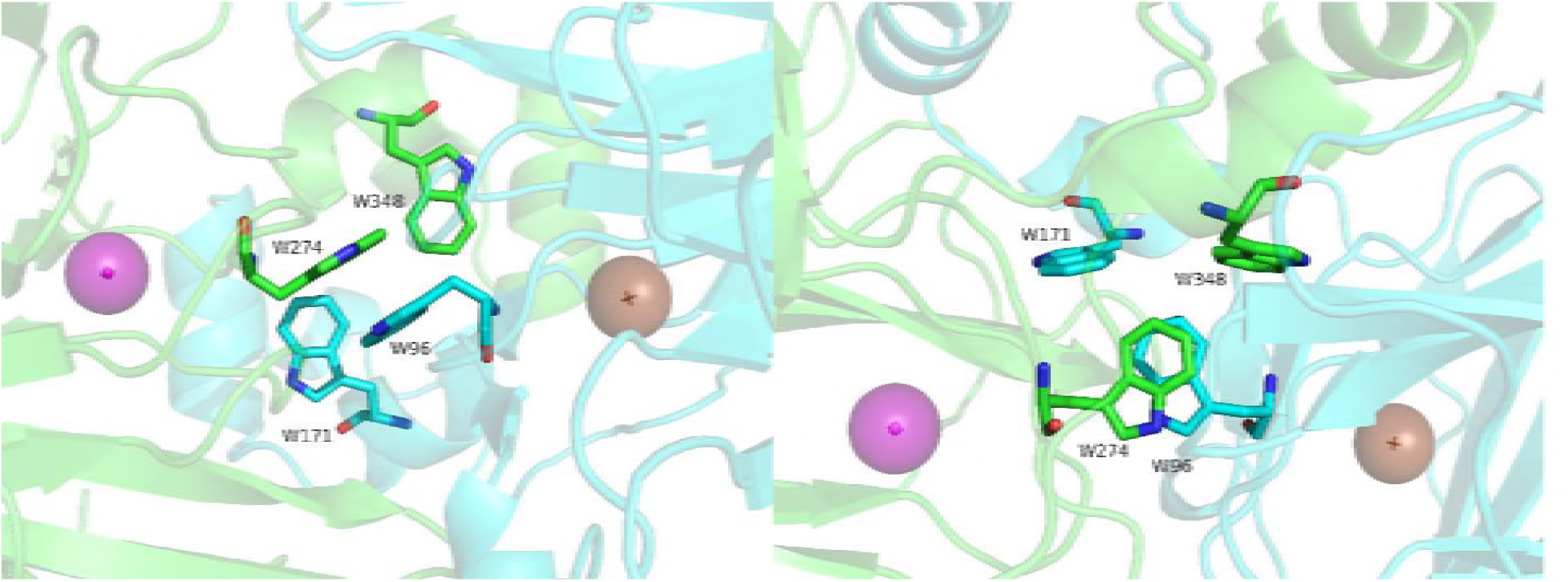
Different views of the four tryptophans making up the hot spot between monomers A (green) and B (cyan). Note that W96 and W274 are π-stacked with an average distance of their molecular planes of about 3.6 Å. W171 and W348 are turned 90 degrees with respect to W96 and W274 and are located near the edges of the π-stacked dimer.

3. Salt Bridges Stabilizing the Quaternary Structure

**Figure S 3:**
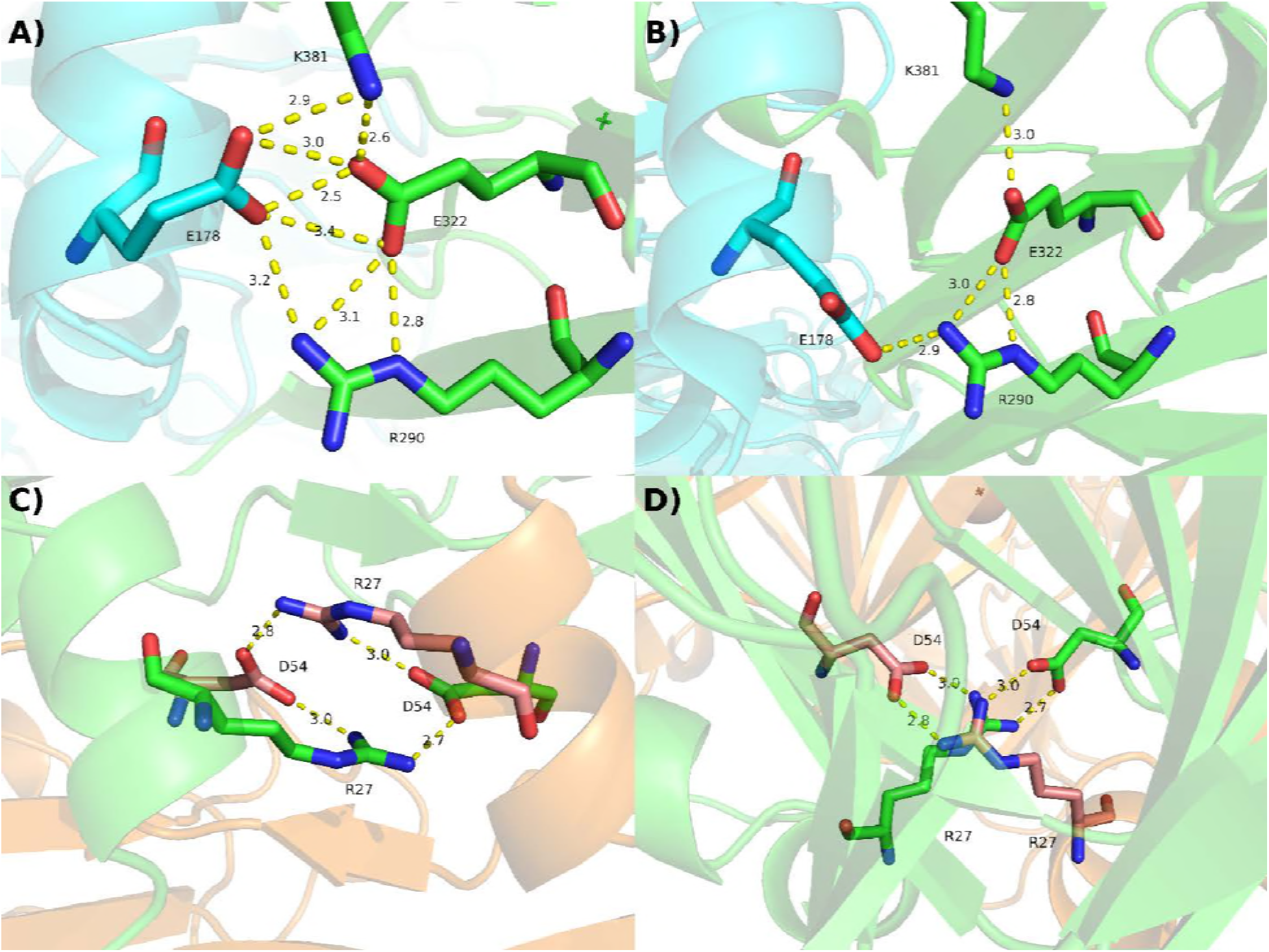
Network salt bridges between adjacent monomers. A) Three residues on monomer A (green) form a networked salt bridge with residue E178 of monomer B near the apex of a triangle. B) same region as in A) but for the high-pH structure 1UW8. C) Quadrangle salt bridge between residues R27 and D54 in monomers A and D. R27 is part of a short α-helical domain I25-Q30 while D54 is part of the strand that connects the bottom parts of the N- and C-terminal cupin domains with each other. D) Same as C) but viewed from the top of monomer A perpendicular to the triangle.

4. Electron density map of the flexible loop region SENST161-165 with the models of both open and closed conformation.

**Figure S 4:**
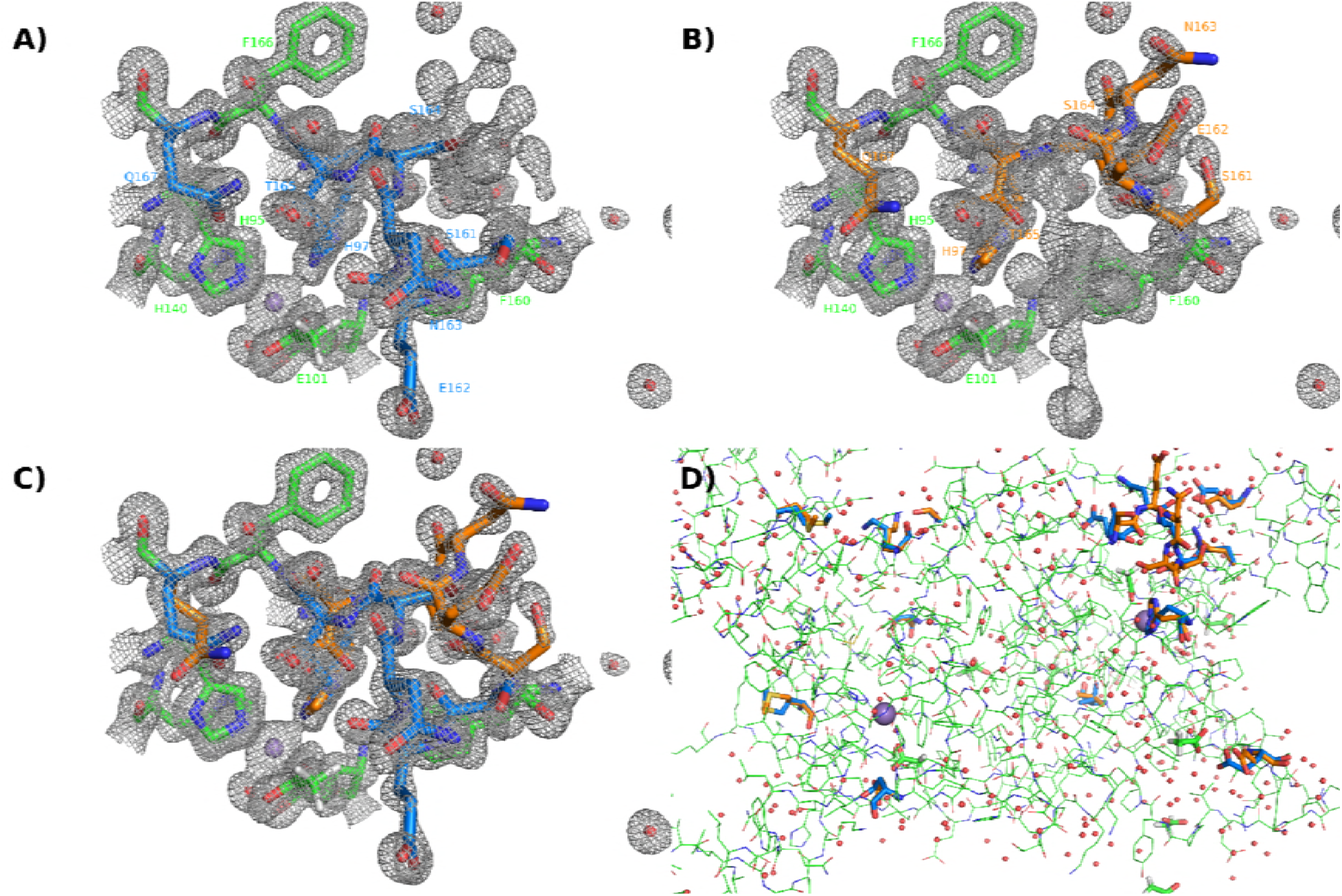
Electron density map (2Fo-Fc, contoured at 1.3 sigma) of the loop region of OxDc that exists in two alternate conformations. The two loops (residues 161-165) were modeled and refined at 60 and 40% occupancy. A) Closed conformation with ‘alt A’ residues colored in ‘marine.’ B) Open conformation with ‘alt B’ residues colored in ‘orange.’ C) Both alternate conformations together. D) Subunit A of OxDC displayed as lines with ‘alt A’ (closed conformation) residues in marine ‘stick’ representation, and ‘alt B’ (open conformation) residues as orange ‘sticks.’

5. Comparison between the closed conformations of 5VG3 (low pH structure) and 3S0M (T165V mutant at high pH).

**Figure S 5:**
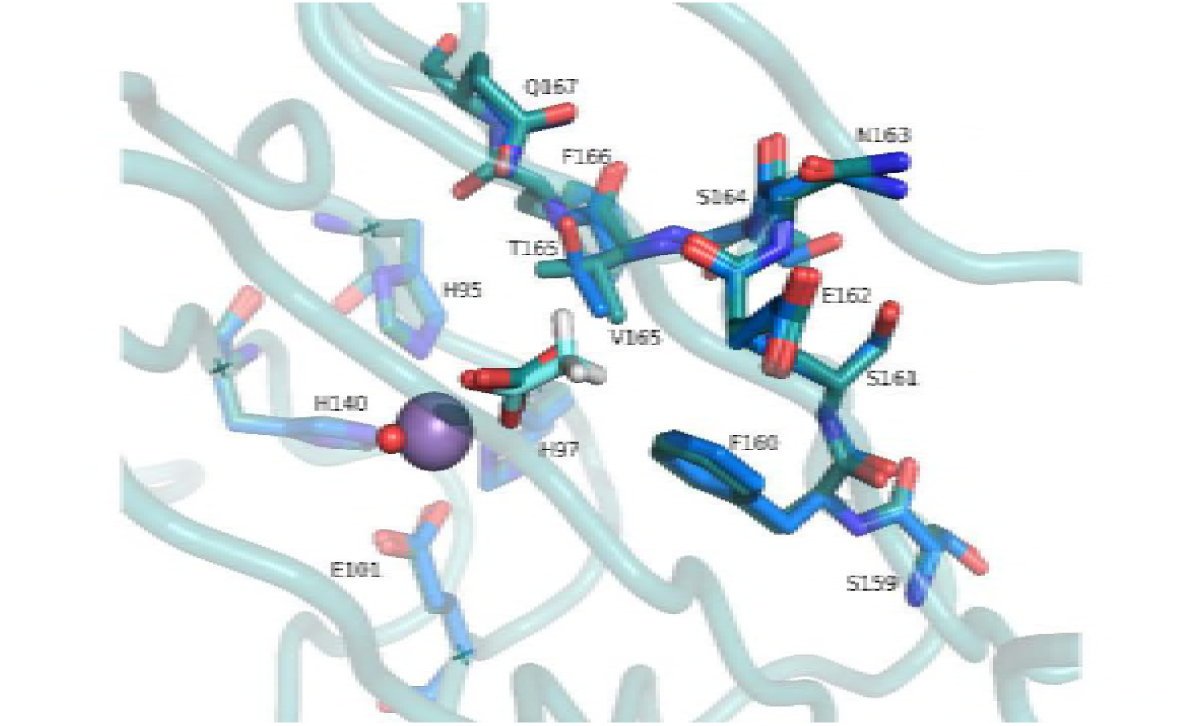
Comparison of the N-terminal binding site of 3s0m (T165V mutant) in teal showing the coordinating residues and the closed loop with adjacent residues, SFSENST(V)FQ159-167. The same residues are shown in blue for the closed conformation (alternate B) of 5vg3. Note the close overlap between the two loop structures. The small molecule ligands acetate (5vg3) and carbonate (3s0m) are also shown. They coordinate to Mn at the same location.

6. B-factors of the flexible loop region.

**Figure S 6:**
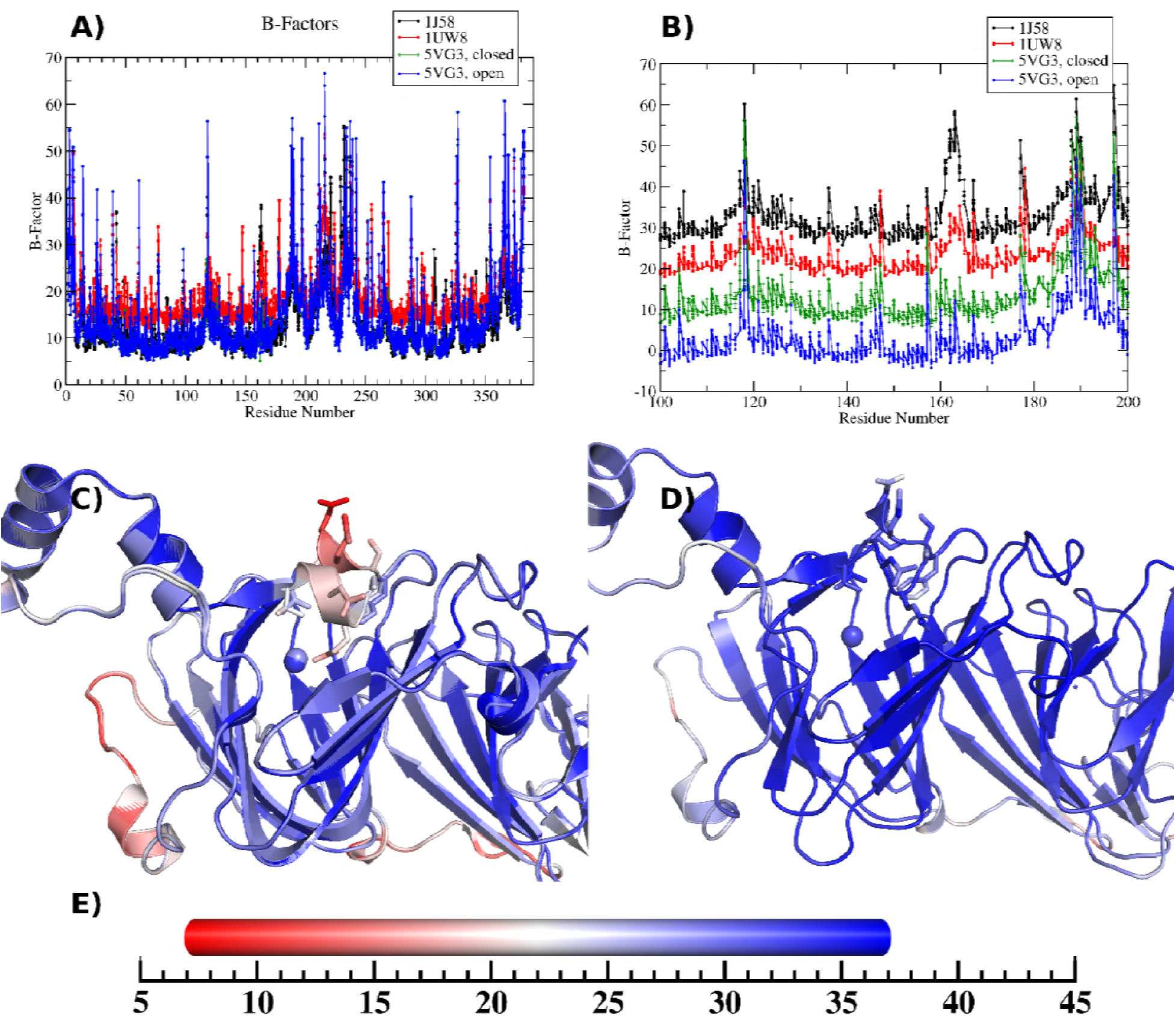
B-factors of the flexible loop region. A) Plot of atomic B-factors for the open (1J58) and closed (1UW8) conformations of OxDC at high pH and those at low pH (5VG3) as a function of residue number. B) Similar to A) but only for residues 100 to 200 and with vertical offsets to better distinguish the different crystal structures. C) and D) are cartoon plots of the high pH (C) and low pH (D) open and closed conformations shown in a color scale (E) that ranges from 7 (red) to 37 (blue) for the atomic B-factors.

7. H-bonding Network around the SENST Loop in the Open and Closed Conformation.

**Figure S 7:**
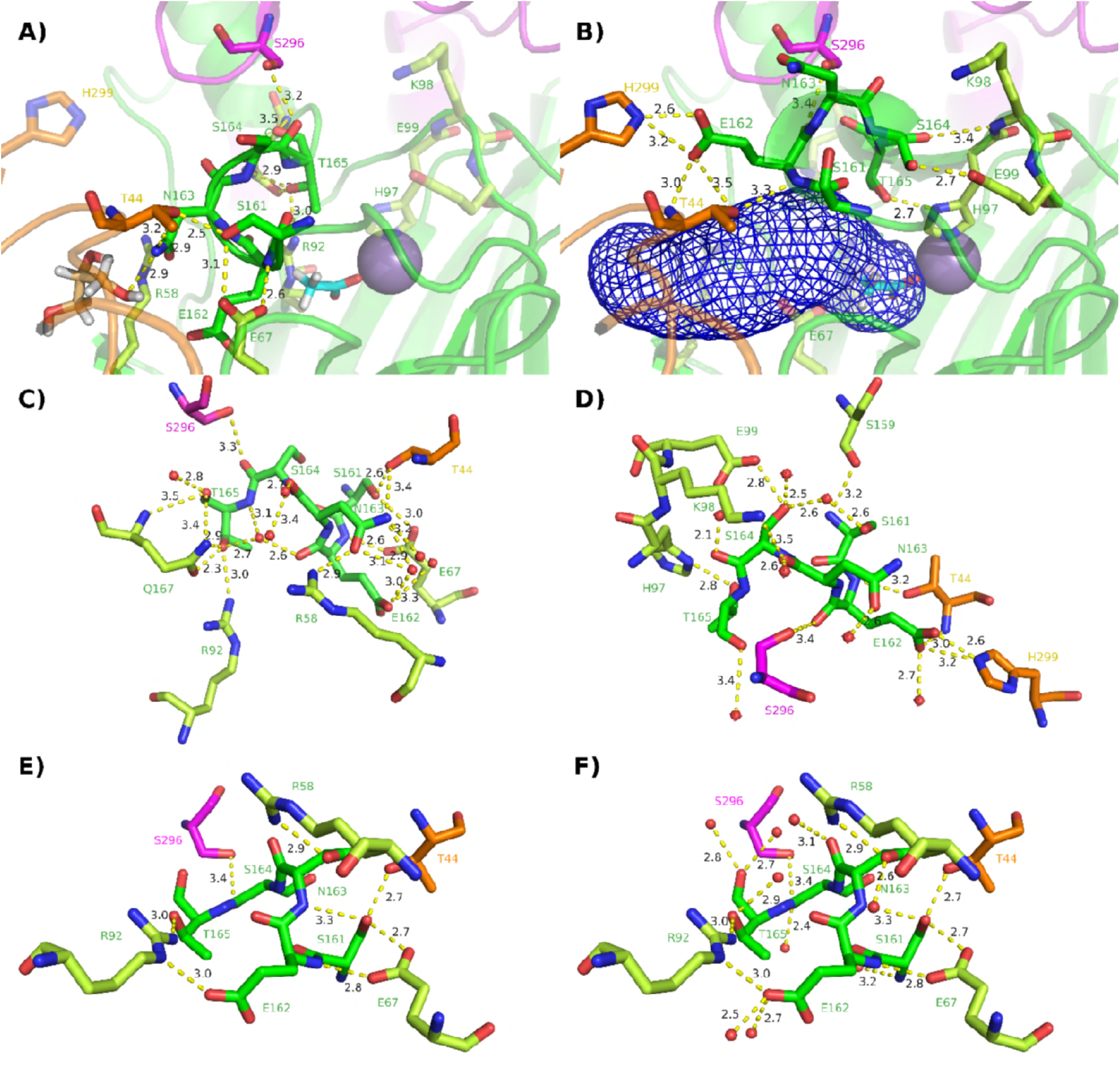
Hydrogen bonding around the SENST loop. A) and C) closed conformation of 5VG3 with and without water (red spheres); B) and D) open conformation of 5VG3 with and without water (red spheres). Color coding: SENST loop green (chain A), amino acids within the same chain limon green, neighboring chain residues magenta (chain C) and orange (chain D). Non-hydrogen bonding residues in a given conformation that show H-bonds to the SENST loop in the other configuration are shown with 50% transparency level. The solvent channel providing access to the N-terminal Mn site in the open conformation (B) was calculated with Caver 3.0.1 and is shown enclosed with a blue mesh. E) and F) H-bonding analysis of the closed conformation at high pH (1UW8) with (F) and without (E) water.

8. Mono- and Bi-dentate Oxalate Binding Models.

**Figure S 8:**
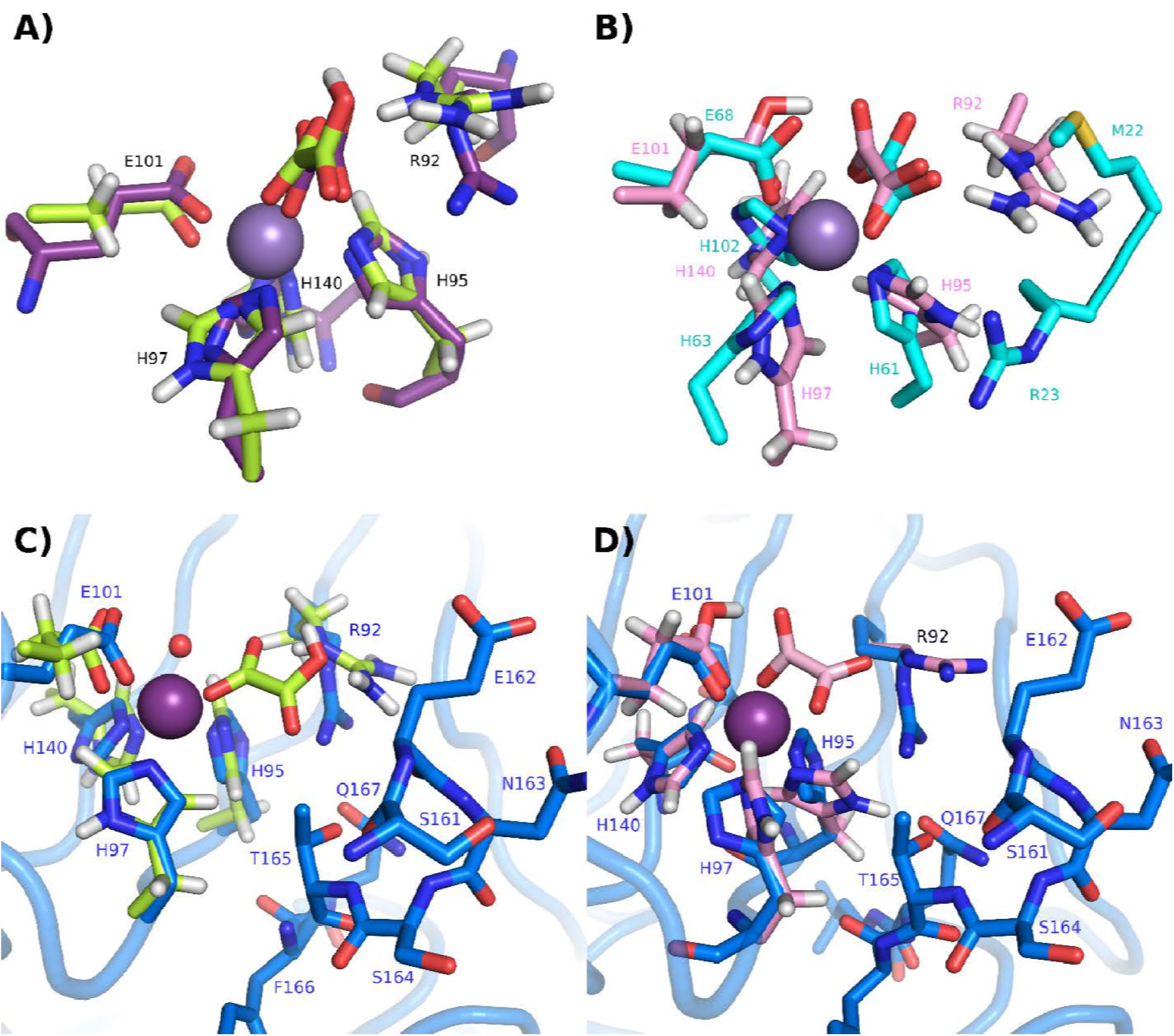
N-terminal binding pocket with oxalate simulated in the calculated mono-dentate (A and C, limon green) and bi-dentate (B and D, pink) coordinations. The simulated oxalate positions are compared to the N-terminal binding site of the E162 deletion mutant, 5HI0 (A, purple), and with the Oxalate in the Mn-site of the putative oxalate decarboxylase tm1287 from Thermotoga maritima (B, cyan, PDB#1O4T). The blue residues in C) and D) represent the low pH crystal structure. 5VG3, in its closed conformation.

9. N-terminal binding pocket with simulated oxalate in mono- and bi-dentate.

**Figure S 9:**
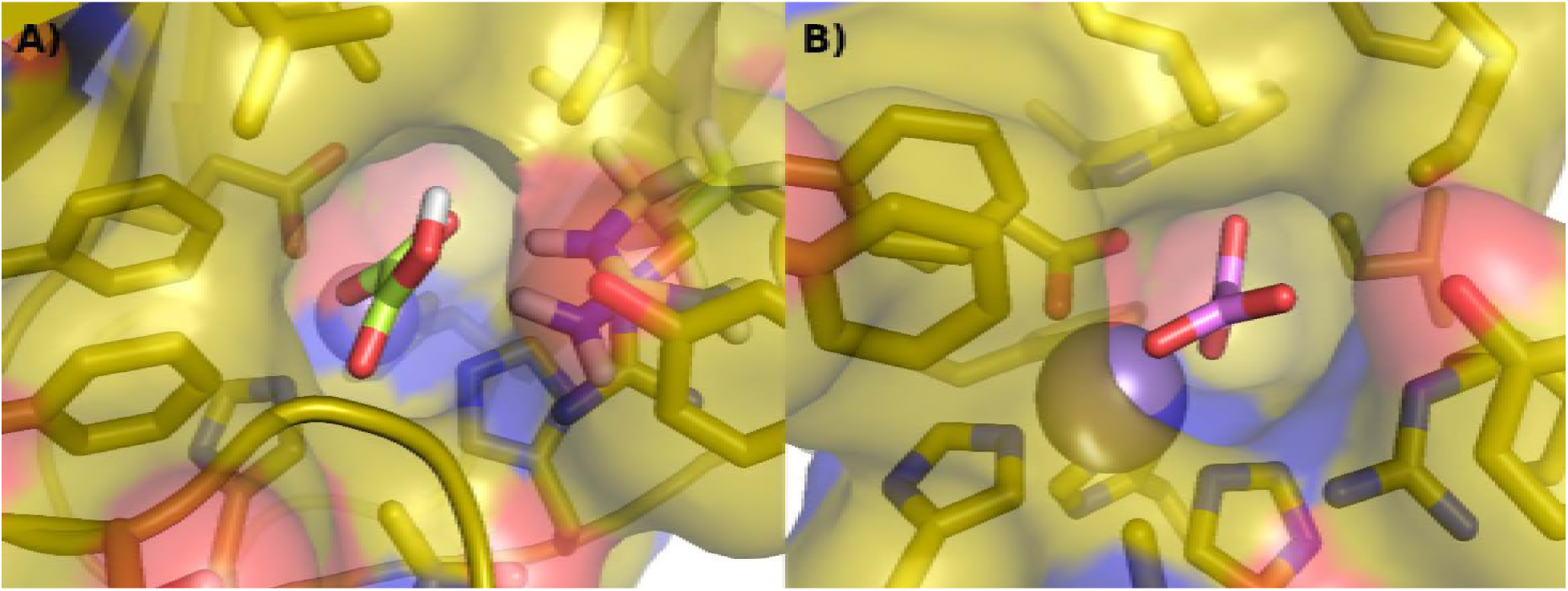
Cavity around the modeled oxalate molecule in the N-terminal binding site of 5VG3 in the open conformation. A) mono-dentate binding mode. B) Bi-dentate binding mode.

10. Protein cavity near C-terminal Mn-binding site.

**Figure S 10:**
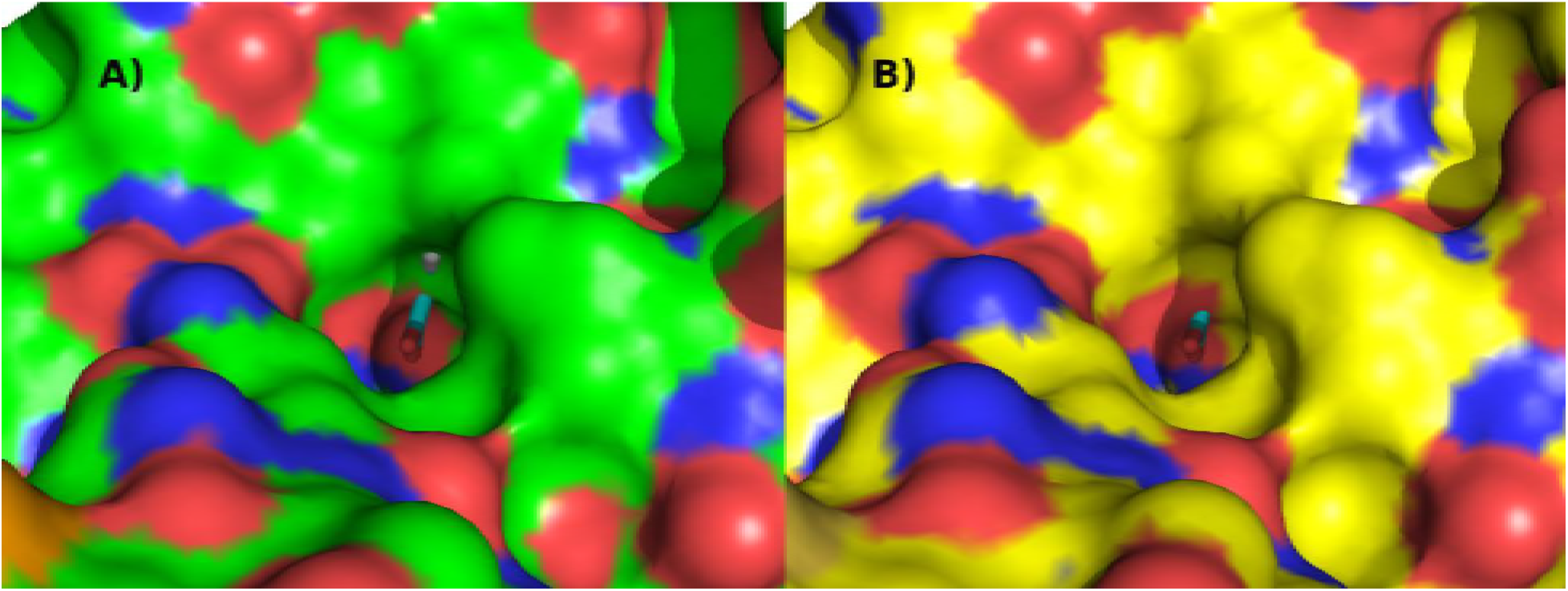
Protein surface near C-terminal site showing the cavity with the acetate molecule from 5VG3 overlaid. One can see that the acetate clashes with part of the protein surface in both high pH structures: A) 1J58 (open loop structure) and B) 1UW8 (closed loop structure).

11. *In-silico* Alanine Scanning Score results for subunit A:

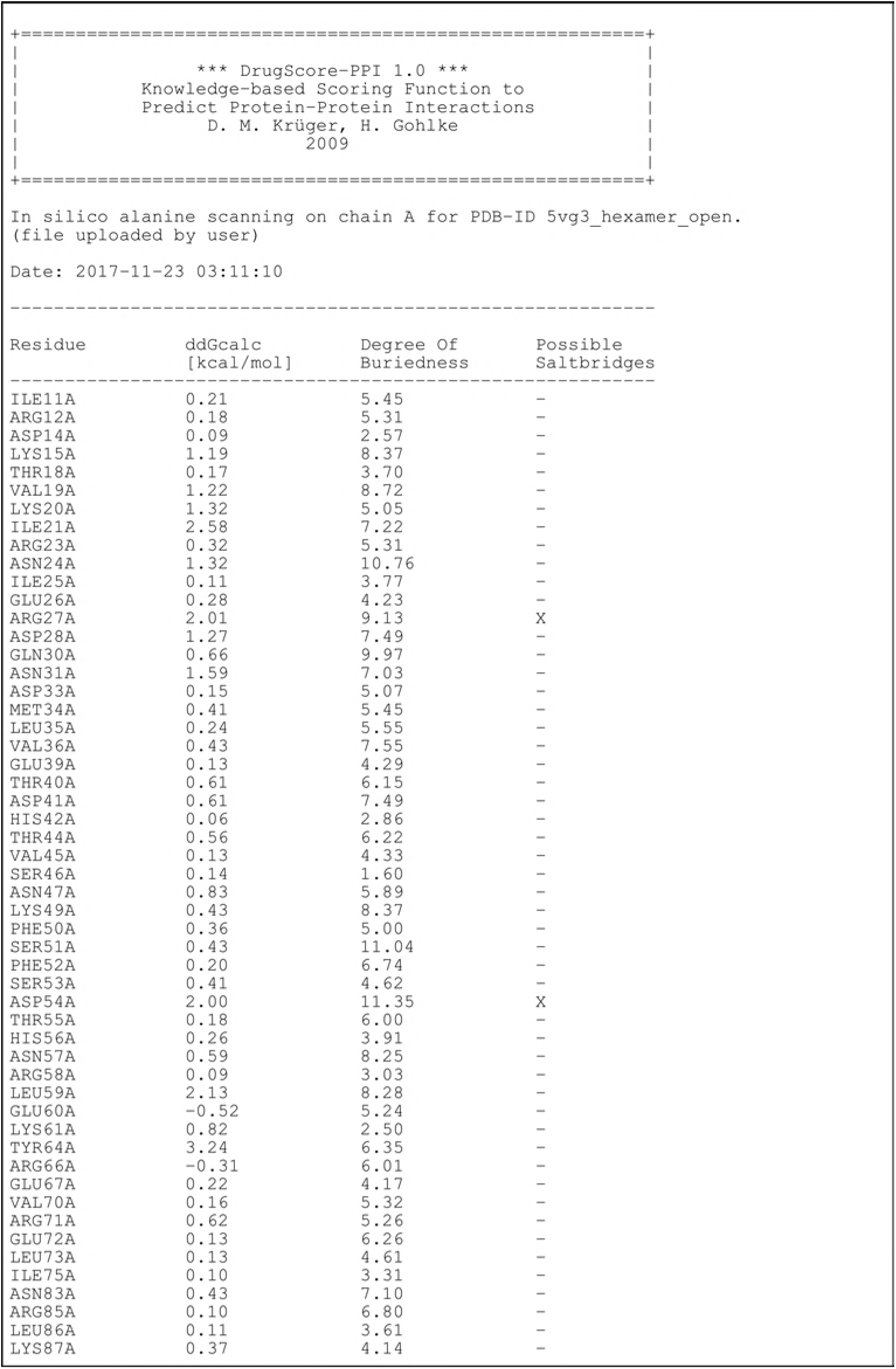

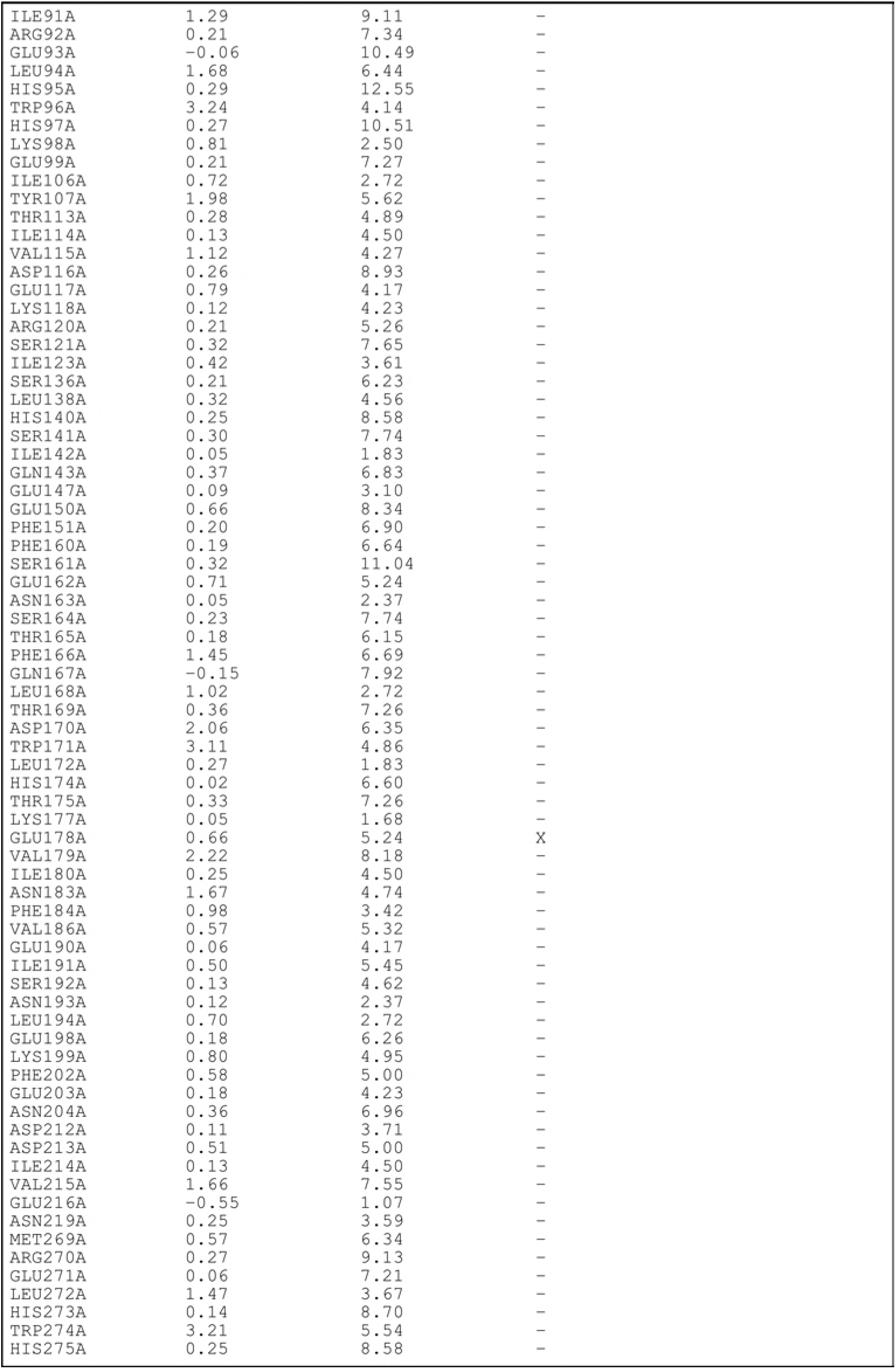

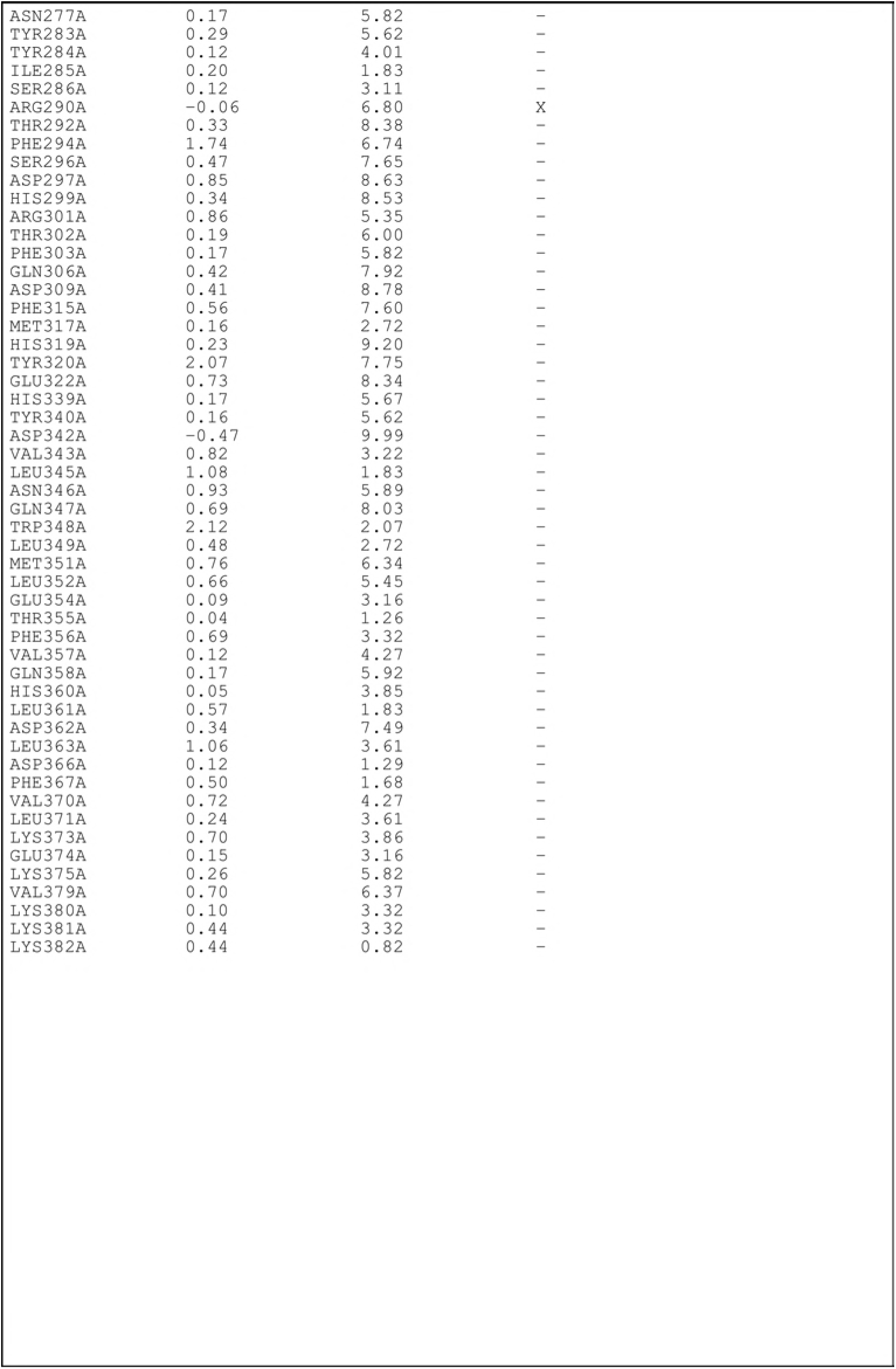

